# PARP knockdown promotes synapse reformation after axon injury

**DOI:** 10.1101/2023.11.03.565562

**Authors:** Micah Y. Belew, Wenjia Huang, Jeremy T. Florman, Mark J. Alkema, Alexandra B. Byrne

**Affiliations:** Department of Neurobiology, NeuroNexus Institute, UMass Chan Medical School, Worcester, Massachusetts, USA

## Abstract

Injured nervous systems are often incapable of self-repairing, resulting in permanent loss of function and disability. To restore function, a severed axon must not only regenerate, but must also reform synapses with target cells. Together, these processes beget functional axon regeneration. Progress has been made towards a mechanistic understanding of axon regeneration. However, the molecular mechanisms that determine whether and how synapses are formed by a regenerated motor axon are not well understood. Using a combination of *in vivo* laser axotomy, genetics, and high-resolution imaging, we find that poly (ADP-ribose) polymerases (PARPs) inhibit synapse reformation in regenerating axons. As a result, regenerated *parp(-)* axons regain more function than regenerated wild-type axons, even though both have reached their target cells. We find that PARPs regulate both axon regeneration and synapse reformation in coordination with proteolytic calpain CLP-4. These results indicate approaches to functionally repair the injured nervous system must specifically target synapse reformation, in addition to other components of the injury response.

## Introduction

After injury, adult nervous systems are often incapable of repairing themselves, resulting in a permanent loss of motor and sensory function. Successful repair of severed axons requires functional axon regeneration, a poorly understood process in which severed axons both regenerate and reform functional synapses with their relevant target cells. Mechanisms that inhibit axon regeneration, particularly in the central nervous system and in the aged peripheral nervous system, have been identified. However, how synapse reformation between a regenerating axon and its target cell is regulated after injury is poorly understood. Identifying the molecular mechanisms that regulate each aspect of functional axon regeneration will fundamentally inform approaches to treat axon injury and disease.

An axon’s ability to regenerate is determined by extrinsic factors such as myelin-associated factors and chondroitin sulfate proteoglycans, and by intrinsic factors such as PARPs, PTEN and DLK-1 (1–4). Known interventions that strongly promote axon regeneration only modestly restore function to the injured CNS (5–7). However, such functional restoration is often the result of sprouting from neighboring intact axons and does not involve synapse formation with the regenerated axons themselves (8, 9). In the peripheral nervous system, where young axons are capable of regeneration, severe damage often inhibits synapse formation or results in inappropriate innervation and chronic pain (10). These outcomes delineate a critical need to understand how synapse reformation can be improved between a regenerated axon and its postsynaptic target.

We previously found that poly (ADP-ribosyl)ation inhibits the first requirement for functional restoration after injury: axon regeneration. Poly (APD-ribose) glycohydrolases function downstream of DLK-1/DLK signaling to positively regulate regeneration of injured *C. elegans* GABA motor axons (11). In opposition to the poly (ADP-ribose) glycohydrolases, poly (ADP-ribose) polymerases (PARPs) inhibit axon regeneration. Poly (ADP-ribosyl)ation is an increasingly studied post-translational modification in which chains of ADP-ribose molecules are transferred from NAD^+^ to substrate proteins. In addition to axon regeneration, the number of biological processes known to be regulated by PARylation is dramatically expanding and includes DNA repair, apoptosis, mitosis, transcription, and chromatin modification (12). Multiple lines of evidence suggest the role of PARPs in axon regeneration is also conserved in mammals. In particular, inhibiting PARPs either genetically or with chemical PARP inhibitors significantly enhances axon regeneration in injured murine cortical neurons *in vitro* (11). In the absence of inhibition, PARylation is upregulated in crushed optic nerves after injury and in cortical neurons placed on substrates that block regeneration, including myelin-associated glycoprotein, Nogo-A, and Chondroitin sulfate proteoglycans. Consequently, inhibiting PARP1 improves neurite outgrowth on inhibitory substrates *in vitro* (13). The role of PARPs in axon regeneration likely extends to human neurons, as exposure to multiple individual chemical PARP inhibitors also inhibits outgrowth of human iPSC derived cortical and motor neurons (14). Currently, redundancy among 18 PARPs and the presence of extrinsic inhibitors of axon regeneration has limited the ability to tease apart the role of PARPs in axon regeneration *in vivo* in a mammalian system (15, 16).

While the finding that PARPs regulate axon regeneration adds to the limited repertoire of genes known to regulate the injury response, the more intriguing observation is that wild-type animals exposed to chemical PARP inhibitors regained more motor neuron function than wild-type animals exposed to DMSO after injuring 15 of the 19 dorso-ventral GABA motor axons that inhibit the body wall muscles of *C. elegans* (11). The observation raises several critical questions, particularly 1) whether the observed functional recovery is caused by an increased ability of individual PARP axons to reform synapses or simply by an increase in the number of axons that regenerate in PARP mutants), 2) whether the enhanced axon regeneration is caused by a loss of PARylation or a parallel increase in NAD^+^ availability, and 3) what are the downstream effectors of functional axon regeneration?

The existence of only two PARP homologs makes *C. elegans* an attractive system to investigate how PARP inhibition regulates the injury response. Beyond the benefit of *C. elegans’* relatively reduced complexity, regulators of axon regeneration can be investigated in a short time frame with single axon resolution in vivo (17, 18), due to its relatively short lifespan, well-characterized nervous system and transparent body (4). Moreover, approximately 60% of *C. elegans* genes are conserved with mammals, including regulators of axon regeneration such as PTEN and DLK (19–23). Together, the enhanced ability to study intrinsic mechanisms of axon regeneration and a tractable number of PARP homologs makes *C. elegans* an informative model to investigate how PARPs regulate the injury response.

Here, we find that inhibiting poly (ADP-ribose) polymerase function improves synapse reformation and restores function to individual injured motor axons. PARPs regulate two arms of functional axon regeneration by inhibiting both axon regeneration and synapse reformation through interactions with the proteolytic calpain *clp-4*, which is homologous to Calpain B in *D. melanogaster* and several human Calpains, including CAPN3, CAPN8 and CAPN9 (24). Calpains have previously been shown to truncate synaptic proteins during development, suggesting the PARPs co-opt this developmental mechanism to inhibit functional regeneration of injured axons in adult animals. Because individual regenerated axons in *parp(-)* and *clp-4(-)* animals reform more synapses than regenerated axons in wild type animals, our results demonstrate that synapse reformation is actively inhibited by PARPs and CLP-4. The parallel finding that developmental synapse formation is grossly wild type in *parp(-)* and *clp-4(-)* animals indicate these genes are capable of regulating synapse formation specifically in the context of the injured or nervous system. Our finding that synapse reformation can be improved relative to wild type regenerated axons demonstrates that synapse reformation is actively inhibited, particularly by PARPs. Therefore, injured wild type axons are prevented from regaining function not only by mechanisms that inhibit axon regeneration, but also by mechanisms that inhibit synapse reformation. Identifying and manipulating genes that inhibit both axon regeneration and synapse reformation is a streamlined and powerful approach to achieving functional repair of the injured nervous system.

## Results

### Loss of PARP function enhances functional axon regeneration of injured axons

We observed that wild type animals exposed to chemical PARP inhibition recover more gross locomotor function than vehicle controls after severing the majority of GABA motor axons that innervate body wall muscles (11). This exciting observation raises the critical question of whether the recovery of motor function is caused by an enhanced ability of individual regenerated *parp(-)* axons to reform synapses with body wall muscles (functional axon regeneration). Alternatively, individual regenerated *parp(-)* axons may simply form the same number of new synapses as wild type axons, however motor function is improved because more axons regenerate in *parp(-)* animals compared to wild type animals. Differentiating between these two possibilities would be highly significant as it could reveal a specific role of PARPs in synapse reformation, and in turn determine whether synapse reformation is specifically inhibited in injured wild type motor axons. Thus, we developed an in vivo approach to measure synapse reformation and functional locomotor recovery with single axon resolution, and asked whether regenerated axons that lack PARPs reformed synapses and regained function more frequently than regenerated wild-type axons.

As chemical PAR inhibition has many off-target effects and does not completely eliminate PAR activity, we generated animals with a complete and specific loss of PARP function. The resulting *parp-1(ok988); parp-2(ok344)* animal contains null mutations in *C. elegans’* only two *parp* genes and is herein referred to as *parp(-)*. The *parp(-)* strain also contains an integrated transgene that expresses cytoplasmic GFP specifically in GABA motor neurons to visualize GABA axons before and after injury (*parp-1(ok988); parp-2(ok344); oxIs12 (Punc-47::GFP + lin-15B*)) (25) (Figure 1A,B). After severing the above axons at the midline using a standard axotomy protocol (18), *parp(-)* animals displayed a robust increase in both the number of axons that initiated regeneration (regeneration, *wt* 59.79% vs. *parp(-)* 84.10%, p=0.001) and the number of axons that regenerated the full width of the animal to reach the dorsal nerve cord (full regeneration, *wt* 20.63% vs. *parp(-)* 41.16%, p=0.001) compared to wild-type animals (Figure 1C,D). The increased ability of regenerating *parp(-)* axons to reach their synaptic partners in the dorsal cord raises the question of whether these individual axons are more or less capable of regaining function compared to individual wild type axons.

**Figure 1.**
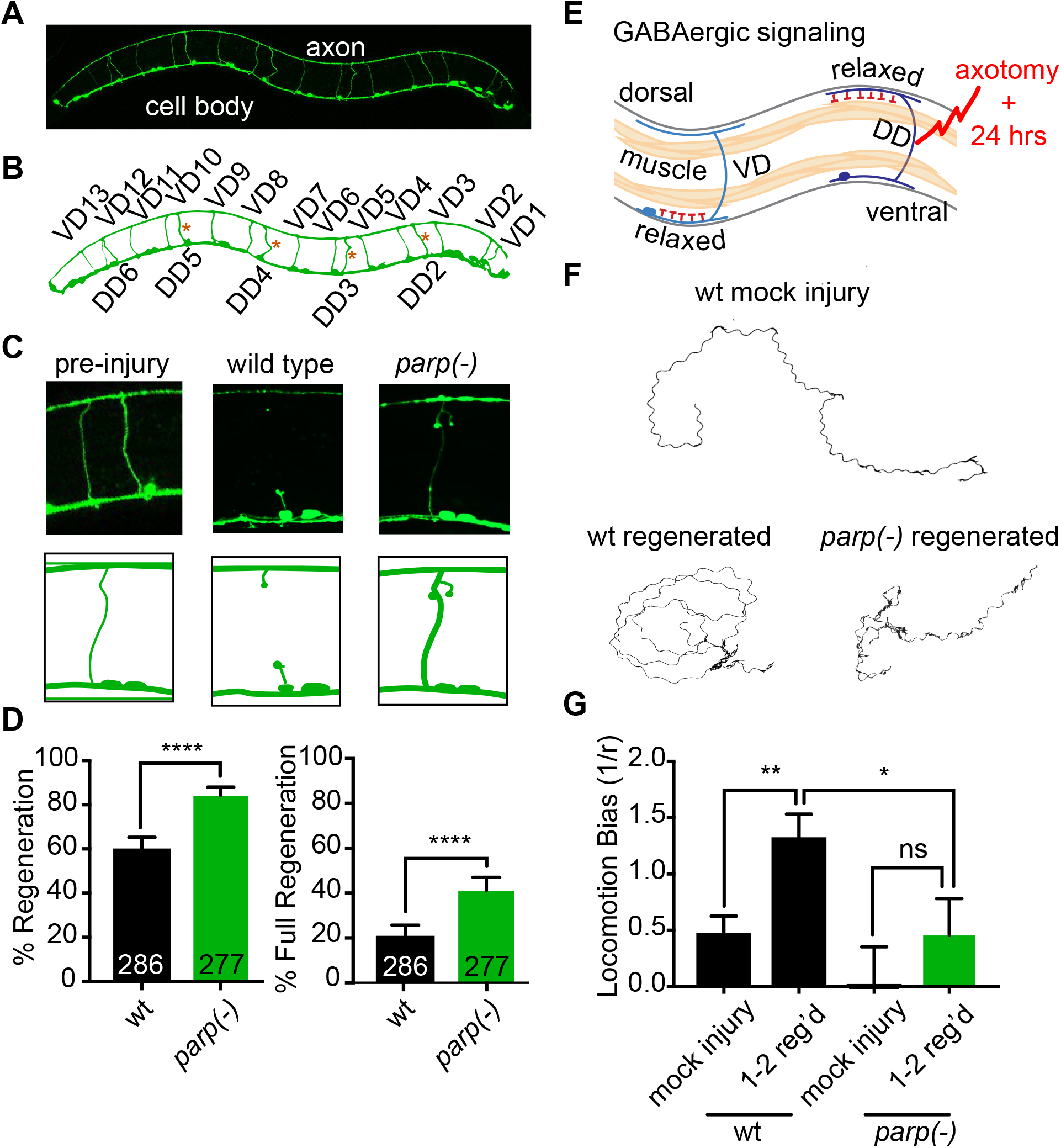
PARP knockdown improves functional axon regeneration. (**A**) The cell bodies of GFP labeled GABA motor neurons lie on the ventral side of the animal and extend axons to the dorsal cord. (**B**) The axons are subdivided into ventrally directed (VD) and dorsally directed (DD), which synapse onto the ventral and dorsal body wall muscles, respectively. (**C**) Severing individual GABA neurons reveals that (**D**) *parp-1(-); parp-2(-)* axons regenerate more frequently and reach their target muscles in the dorsal cord (full regeneration) more often than wild type axons. Axon regeneration was calculated by determining the percentage of injured that initiated a growth cone or fully regenerated 24h post-injury. Error bars indicate 95% confidence intervals. Fisher’s exact test was used to determine significance. (**E**) The GABA motor neurons relax their target muscle cells in opposition to cholinergic axons which innervate the muscle on the other side of the animal. This dichotomous signaling pattern creates a sinusoidal locomotion. (**F**) Animal movement was recorded 24 hours after severing axons DD2-5 with an automated Worm Tracker. Animals that underwent a mock injury maintain a wild type sinusoidal pattern of movement. However, injured wild type animals that regenerate 1-2 axons are unable to relax their dorsal body wall muscles and exhibit a strong dorsally directed locomotion bias. In contrast, *parp(-)* mutants that regenerate the same number of axons recover a relatively wild type locomotion pattern compared to wild type animals. (**G**) The locomotion bias was calculated from the curvature of the animal’s tracks and indicates *parp(-)* mutants recover more dorsal body wall function than wild type animals (see Methods). Error bars indicate standard error of the mean. Significance was determined with a student’s t-test.

Determining whether individual axons undergo functional axon regeneration requires a highly sensitive assay to quantify neuronal function. Dorsally directed (DD) and ventrally directed (VD) GABA motor neuron activation is required to alternately relax the dorsal and ventral body-wall muscles (Figure 1B;E, blue). Coordination of GABAergic activity with cholinergic axons, which when activated induce contraction of body-wall muscles on the opposite side of the animal, gives rise to a behavior consisting of a highly symmetric sinusoidal pattern of movement. We hypothesized that severing four DD-type GABA neurons (DD2-5) would disrupt the animals’ ability to relax the relevant dorsal body-wall muscles, resulting in a quantifiable dorsally directed locomotion direction bias that might be reversed by functional regeneration. We used an automated worm tracker to compare locomotor behavior between wild-type and *parp(-)* animals in which a small and comparable number of injured GABA motor axons had fully regenerated (26). When four of the dorsally-innervating DD-type GABA neurons were axotomized, wild-type animals lost the ability to relax the relevant dorsal body-wall muscles, inducing a bend of the animal in the ventral direction. As a consequence, injured animals displayed a circular ambulatory pattern, as a result of a pronounced dorsally directed locomotion bias 24 hours after injury (Figure 1F,G). In contrast, *parp(-)* animals displayed a reduction in dorsally directed locomotion bias after the same amount of time after injury (*wt* 1.32 vs. *parp(-)* 0.46, p=0.0427) (Figure 1F,G). Importantly, the difference in locomotor recovery was observed between wild-type and *parp(-)* animals when the analysis was restricted to animals that had fully regenerated the same number of axons. These data suggest the difference in functional recovery between *parp(-)* and wild-type animals is unlikely to be caused by an increase in number of axons that regenerate in *parp(-)* animals. Rather, it is likely to result from an enhanced ability of individual regenerated *parp(-)* axons to restore function to the neuromuscular junction. These results raise the significant possibility that in addition to inhibiting axon regeneration, PARPs have a specific role in limiting synapse formation.

### Loss of PARP function induces synapse reformation

The genetic and cellular mechanisms that determine whether a regenerated axon forms synaptic connections with its target cells are crucial for functional regeneration but are poorly understood. We asked whether wild type animals were unable to regain GABA motor neuron function because the regenerated axons were incapable of reforming synapses. *C. elegans* motor axons from en passant synapses with the body wall muscles. Their synapses are composed almost exclusively of evolutionarily conserved synaptic components, which are transported from the cell body to the synapse during development. Whether regenerating wild type motor axons are capable of reforming synapses is not known. Therefore, we compared the number of GFP-labeled SNB-1/synaptobrevin puncta, which labels synaptic vesicles, before and after injuring a single DD6 axon (Figure 1B and Supplemental Figure 1), and noted whether the axon did or did not fully regenerate to the mCherry labeled dorsal cord (FR(+) and FR(-), respectively) (Figure 2A,B,C). As previously reported, the severed axon fragments distal to the site of injury persisted 24 hours after injury (Figure 2B (27). However, there were significantly fewer synaptic puncta in these distal fragments compared to intact axons that had not been injured (FR(-) vs mock injury) (Figure 2D,F). In parallel, the total fluorescent intensity of synapses was also significantly smaller than before injury (Supplemental Figure 2A). These data suggest that synapses formed during development had deteriorated after injury.

**Figure 2.**
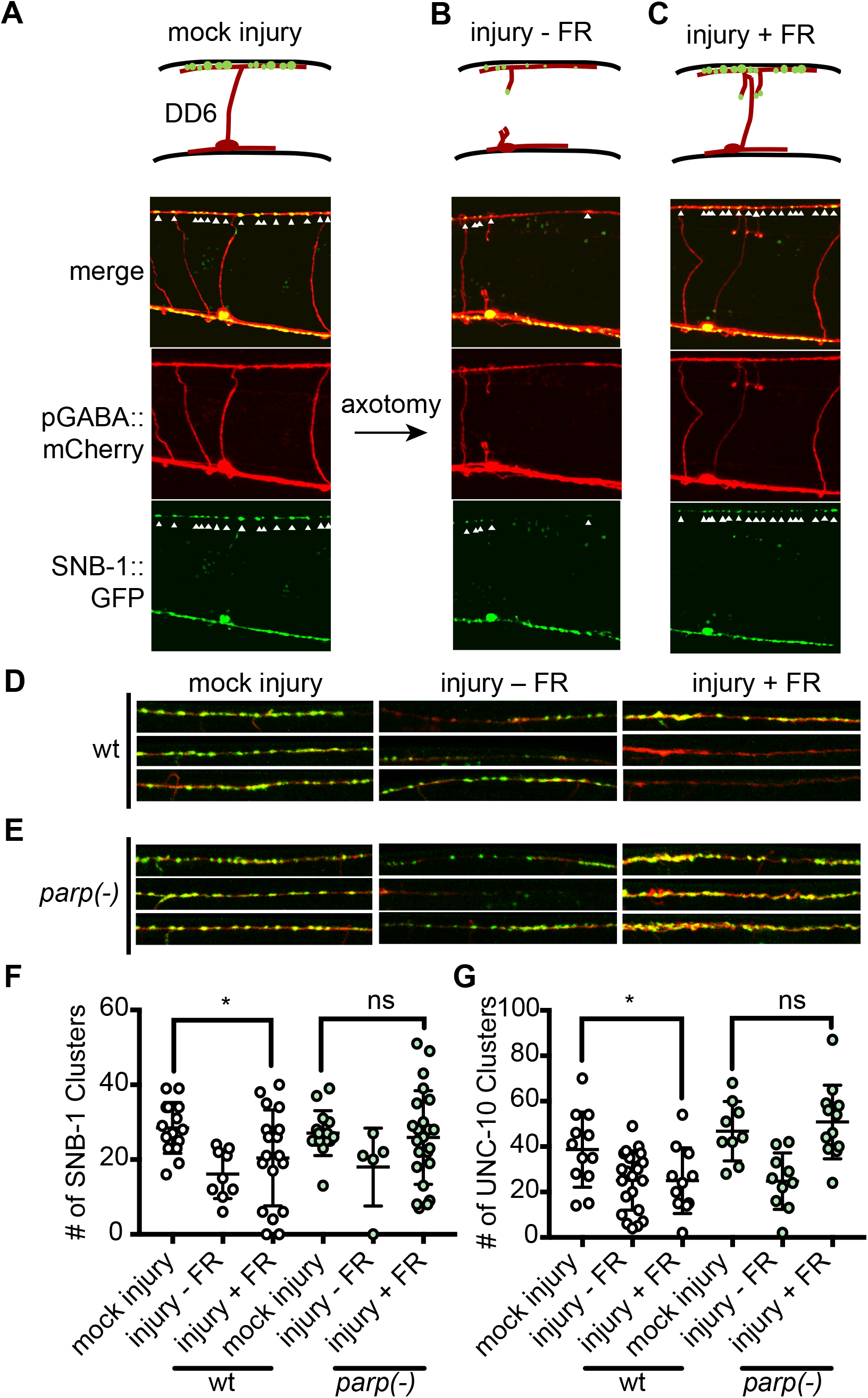
PARP knockdown inhibits synapse reformation within individual regenerated axons. (**A**) The synapse reformation phenotype was measured in mock injured, (B) not fully regenerated (FR(-)) and (C) fully regenerated (FR(+)) DD6 axons 24h-30h post-injury. Axons express *unc-47p::mCherry* and arrows indicate clusters of synaptic SNB-1::GFP. Error bars represent standard deviation. t-test was used to determine significance. (**D**) Representative images of synaptic vesicles in wild type DD6 axons that were uninjured, not fully regenerated, or fully regenerated. (**E**) Representative images of synaptic vesicles in wild type *parp(-)* DD6 axons that were uninjured, not fully regenerated, or fully regenerated. (**F**) Quantification of synaptic SNB-1 clusters in wild type and *parp(-)* animals. (**F**) Quantification of synaptic UNC-10 clusters in wild type and *parp(-)* animals. Each dot represents the number of puncta in one animal. For each genotype, mock injury, injury - FR, and injury + FR were compared. *P < 0.05, **P < 0.01, ***P < 0.001, ****P < 0.0001.

We next asked whether the apparent loss of synaptic puncta was indeed an actively regulated process. The conserved dual leucine zipper MAPKKK DLK-1/DLK/LZK/Wnd is capable of destabilizing developmental synapse formation in *C. elegans* and *D. melanogaster* (20, 28, 29). In wild type animals, DLK-1 function is inhibited during developmental motor neuron presynapse formation by the E3 ubiquitin ligase RPM- 1/PAM/Highwire (28, 30, 31). In the absence of RPM-1, DLK-1 stabilizes *cebp-1* mRNA, which in turn disrupts synapse development (20). We asked whether DLK-1 function was responsible for the observed loss of synaptic puncta after axonal injury. Since DLK-1 is held in an ‘off state’ during development, loss of DLK-1 function does not impede developmental synapse formation (28). We found that similarly, the number of synaptobrevin puncta was not affected by loss of DLK-1 function in uninjured axons. However, synapse destabilization was dependent on DLK-1 in FR(-) axons. The number of synaptic vesicles in FR(+) axons could not be determined because DLK-1 is activated by injury and required for motor axon regeneration (19, 20).Therefore, while DLK-1 is inhibited from disrupting synapse development, its activation upon injury allows it to destabilize synapses after injury (Supplemental Figure 2C, D), Together, these data indicate that the observed synapse destabilization is likely an actively regulated process.

The inability of wild type animals to regain function (Figure 1F,G) prompts the hypothesis that regenerated GABA axons are incapable of reforming functional synapses with the body wall muscles. We found that even when wild type axons had fully regenerated, the number of synaptic puncta was not significantly different from FR(-) axons (Figure 2D,F). Consistently, they also had significantly less synaptic puncta than before axotomy (Figure 2D,F and Supplemental Figure 2A). Therefore, complete axon regeneration is not sufficient to restore synaptic puncta to wild type axons.

To eliminate the possibility that the above observations related just to the distribution of presynaptic vesicles, and not to bona fide synapses, we also investigated how the active zone protein UNC-10/RIM is regulated after injury. We found that UNC-10 puncta were lost in axons that did not regenerate to the dorsal cord and were not restored to pre-injury numbers in wild type axons that had regenerated to the dorsal cord (Figure 2G and Supplemental Figure 2B). Together, in agreement with the finding that regenerated wild type axons did not regain function, the data indicate synapse reformation is disrupted in wild type GABA motor axons, even when axons have reached their pre-injury synaptic partners.

The finding that *parp(-)* animals partially recover normal locomotion compared to wild type animals after axon injury raises the possibility that *parp(-)* mutant animals regulate functional recovery by inhibiting synaptic reformation. We asked if and to what extent synapses are restored after injury when axons have fully regenerated (FR(+)) to the dorsal cord in *parp(-)* mutant animals. We found that like wild type, *parp(-)* FR(-) axons had fewer synaptic punctae compared to mock uninjured axons, suggesting PARP function does not regulate degeneration of the synaptic puncta or promote DLK-1 function. In contrast, individual fully regenerated *parp(-)* axons had similar numbers of synaptic vesicles and active zones as mock injured *parp(-)* axons. This was true when we quantified the number of fluorescently labeled synaptobrevin puncta (SNB-1::GFP) (*parp(-)* 25 vs 26, p>0.05;) and the number of active zone protein UNC-10/RIM punctae (*parp(-)* 47 vs 44, p>0.05;) (Figure 1F,G and Supplemental Figure 2A,B). Therefore, compared to wild type axons, loss of PARP function improves synapse reformation in regenerated axons. A potential explanation for the above observations is that the enhanced regenerative ability of PARP axons induces their earlier arrival at the dorsal cord compared to wild type axons, providing more time to form synapses. However, the ratio of fully regenerated axons to axons that initiated axon regeneration (FR(-)/FR(+)) is the same in wild type and *parp(-)* axons, indicating they regenerate at the same speed (Figure 1D). Therefore, our data provides compelling evidence that PARPs inhibit synapse formation in regenerated axons.

In sum, the observed restoration of function, presynaptic vesicles and active zones, specifically in fully regenerated *parp(-)* axons, indicates that disrupting PARP function restores functional axon regeneration by regulating synapse formation independently from its role in axon regeneration

### PARPs function intrinsically to regulate axon regeneration

To determine the mechanisms by which PARPs inhibit functional repair, we began by investigating how PARPs inhibit axon regeneration. We first asked where PARP function is required to regulate axon regeneration. To do so, we knocked down *parp-1* and *parp-2* specifically in GABA neurons by feeding RNAi to animals in which RNAi is restricted to the GABA nervous system (32–34). Consistent with previously reported rates of regeneration in wild type animals, we found 65.48% of injured axons regenerated when animals were fed *E. coli* containing the empty RNAi plasmid, L4440. In contrast, 88.06%, 83.87%, and 84.0% of injured axons regenerated in animals fed RNAi targeting *parp-1, parp-2*, or both *parp-1* and *parp-2* RNAi, respectively (Figure 3A and Supplemental Figure 3A). In all three assays, the frequency of regeneration in animals lacking *parp* function in GABA neurons was significantly higher than controls. Therefore, PARP function is required in GABA motor neurons to inhibit axon regeneration.

**Figure 3.**
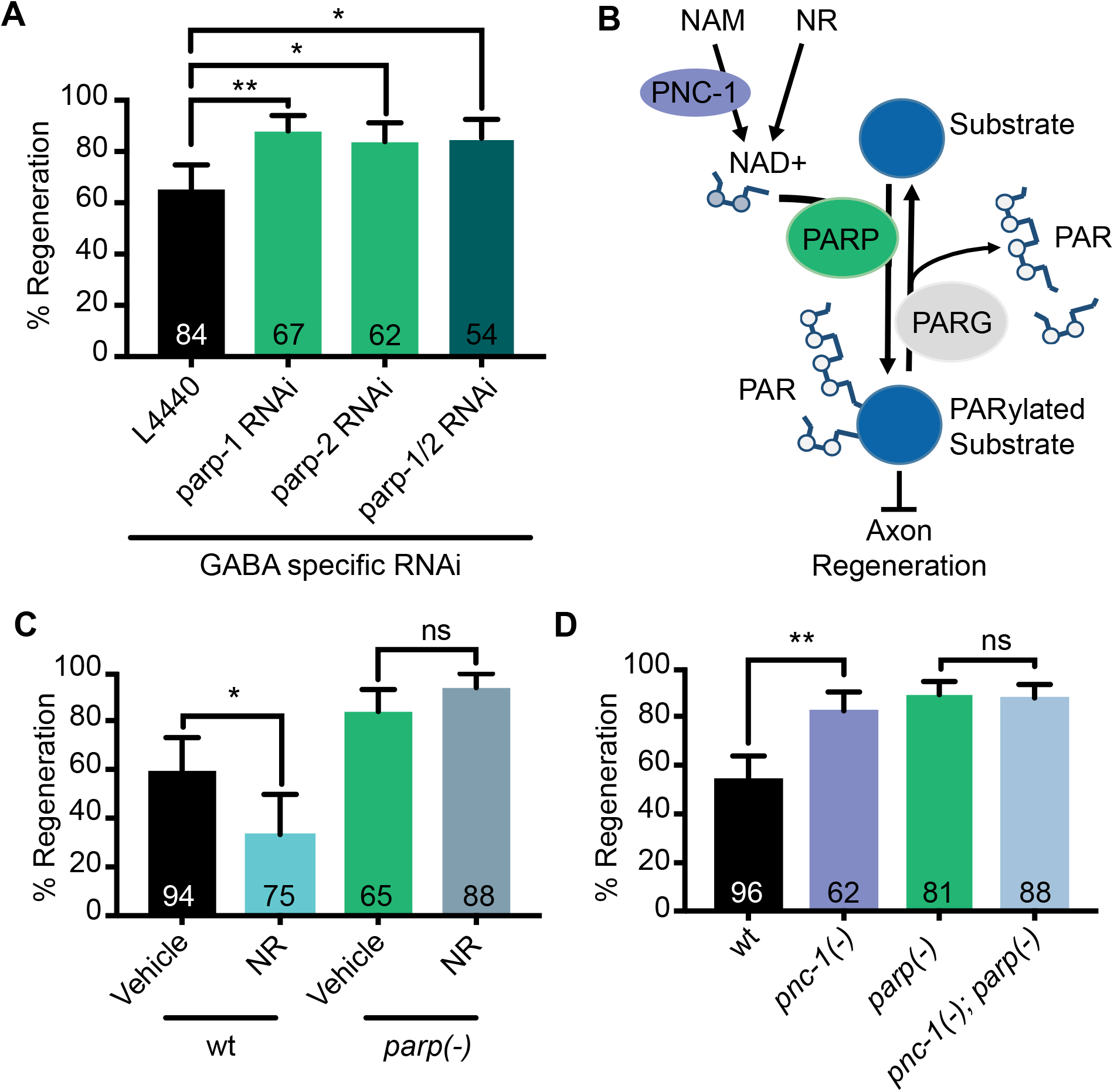
PARP genes regulate axon regeneration cell-autonomously by PARylating substrates. (**A**) Axon regeneration was quantified in animals in which RNAi is restricted to GABA neurons (see Methods). Animals were fed negative control L4440, *parp-1*, *parp-2*, or *parp-1 and parp-2* RNAi. RNAi targeting *parp-1*, *parp-2*, or both *parp-1* and *parp-2* increased axon regeneration relative to L4440. (**B**) Proposed model for the role of PARylation in axon regeneration: PARPs consume NAD^+^ to add poly (ADP-ribose) (PAR) to substrate proteins and inhibit axon regeneration, whereas removal of PAR from substrates by PARGs promotes axon regeneration. (**C**) Axon regeneration was measured in vehicle (water) and 1.25mM Nictinomide riboside (NR) treated animals. NR inhibits axon regeneration in *wildtype* animals, which is rescued by loss of *parp* function, indicating *parp* functions downstream of NR to inhibit axon regeneration. (**D**) The role of *pnc-1* in axon regeneration was measured in *wildtype* and *parp(-)* mutant animals. The *pnc-1(-)* mutation enhanced axon regeneration in wild type animals but not in *parp(-)* mutants, indicating a lack of NAD^+^ did not exacerbate the *parp*(-) phenotype. Error bars indicate 95% confidence intervals. Fisher’s exact test was used to determine significance. *P < 0.05, **P < 0.01, ***P < 0.001, ****P < 0.0001.

### PARPs do not inhibit axon regeneration by increasing NAD^+^ availability

PARPs transfer ADP-ribose to proteins using NAD^+^ as a substrate, a process called PARylation. In addition to decreased PARylation of substrate proteins, a well characterized consequence of inhibiting PARP function is accumulation of NAD^+^ (Figure 3B) (35). The increase in NAD^+^ presents the possibility that in the absence of PARP function, either decreased PARylation of substrate proteins or accumulation of NAD^+^ enhances axon regeneration. We hypothesized that if NAD^+^ accumulation is responsible for the observed increase in axon regeneration in *parp(-)* animals, adding the exogenous precursor of NAD^+^, nicotinamide riboside (NR), would also increase axon regeneration. Instead, we found that NR supplementation decreased axon regeneration relative to animals fed vehicle controls (NR 33.33% vs vehicle 59.57%, p=0.001) (Figure 3C). These results are consistent with the finding that the nicotinamide mononucleotide adenylyltransferase enzyme NMNAT-2 inhibits regeneration of PLM mechanosensory axons in *C. elegans* (36), and suggest that the increased regeneration seen in *parp(-)* animals is not caused by increased NAD^+^. An alternative possibility is that PARylation of substrate proteins regulates axon regeneration. We reasoned that if so, increased NAD^+^ availability might enhance PARP function, thereby further inhibiting axon regeneration, as seen when NR was added to the animals’ diet. To test this possibility, we asked whether loss of PARP function would suppress the increased regeneration phenotype caused by NR supplementation. Indeed, we found the NR-mediated decrease in axon regeneration was suppressed in *parp(-)* animals (83.78% vs. 93.55%, p>0.05) (Figure 3C). To further investigate how NAD^+^ regulates axon regeneration, we quantified axon regeneration in animals with compromised NAD^+^ synthesis. *C. elegans* require the gene *pnc-1 (Pyrazinamidase and NiCotinamidase-1)* to salvage nicotinamide and synthesize NAD^+^ (Figure 3B) (35). Consistent with the decreased axon regeneration observed upon NAD^+^ supplementation, axon regeneration was increased in *pnc-1*(-) animals compared to wild type controls (*pnc-1* 82.26% vs *wt* 54.17%, p=0.0003) and this increase was not enhanced in animals that also lacked *parp* function (88.89% vs 87.505, p>0.05) (Figure 3D). Together with our previous finding that PARGs, which remove poly(ADP-ribose) from substrates, are required for axon regeneration (11), these results suggest that PARylation, and not NAD^+^ accumulation, is responsible for the increased axon regeneration observed in *parp(-)* animals (Figure 3B).

### *clp-4* functions with PARPs to inhibit axon regeneration

To determine the genes that mediate PARP function in injured axons, we performed a candidate screen of previously reported PARP substrates. We hypothesized that if a particular candidate functioned with PARPs to regulate axon regeneration, they would themselves have an axon regeneration phenotype when knocked down. The majority of tested candidates interact with PARP to regulate one of four well-described cellular processes: the DNA damage response, the immune response, cell necrosis, or nucleotide biogenesis (37–42).

Cellular necrosis is regulated by an interaction between PARPs and Calpains, which are calcium dependent proteolytic enzymes that selectively cleave proteins. Cleavage can either activate their targets or prime them for degradation. This prompted us to investigate whether three of the *C. elegans* calpains (*clp-1*, *clp-4*, and *clp-7*), which are expressed in GABA motor neurons (43), regulate axon regeneration. We found that injured axons in animals with a null loss of function mutation in *clp-4* regenerated significantly more frequently than injured wild type axons (*clp-4(-)* 81.67% vs *wt* 62.30%, p=0.0255) (Figure 4A,B,C). However, the number of injured axons that reached the dorsal cord was the same in wild type and *clp-4(-)* animals, indicating the calpain inhibits growth cone initiation but does not inhibit extension (Figure 4F). To determine whether CLP-4 and the PARPs function in the same genetic pathway to inhibit regeneration, we built a *parp(-);clp-4(-)* mutant and asked whether regeneration was further inhibited in the compound mutant compared to *parp(-)* and *clp-4(-)* animals. Loss of *clp-4* did not alter the frequency of growth cone formation or extension of *parp(-)* animals (Figure 4E,G). Therefore, PARPs and *clp-4* function together to inhibit axon regeneration.

**Figure 4.**
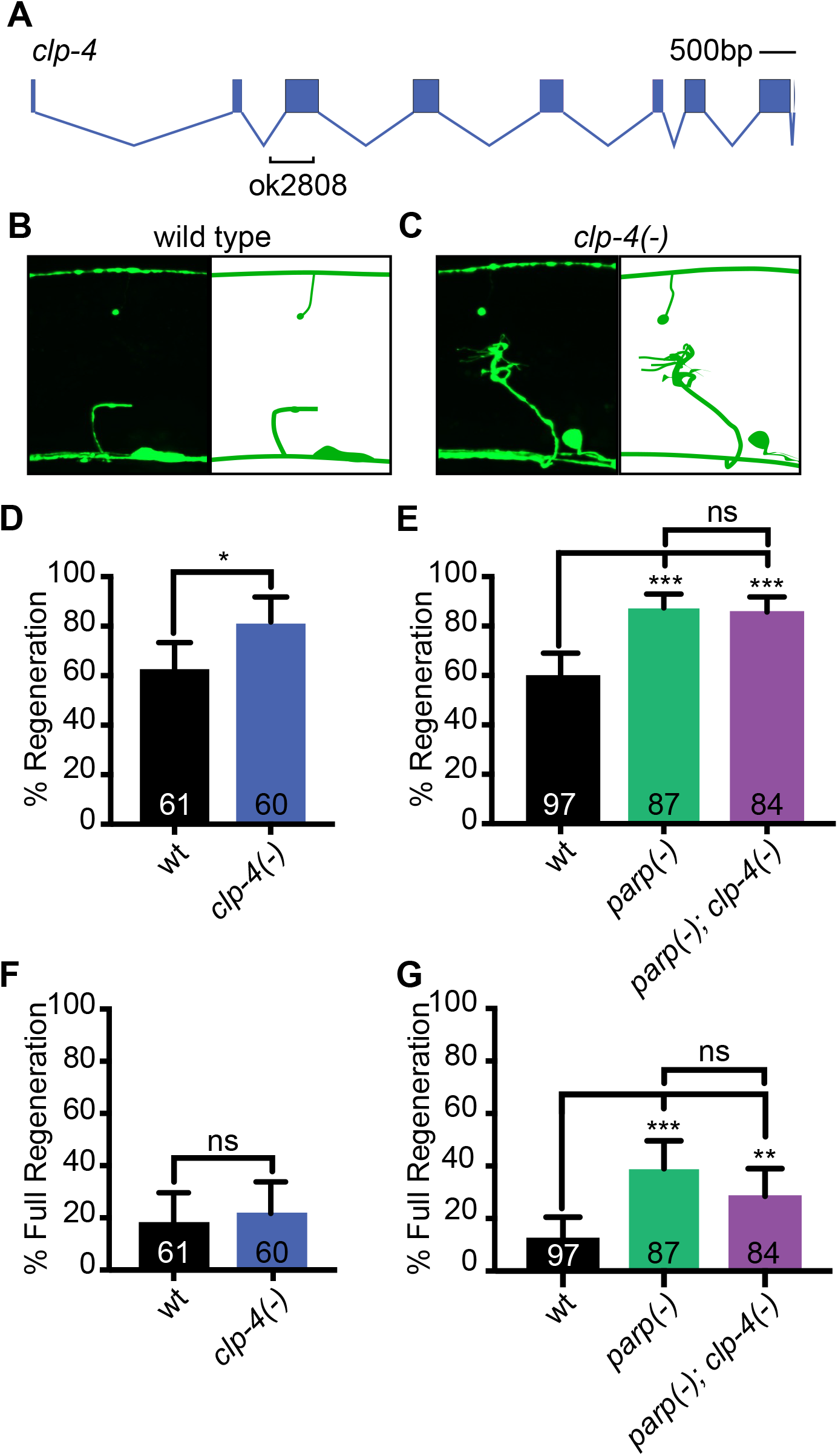
CLP-4/Calpain regulates axon regeneration with PARP genes. (**A**) The ok2808 allele induces an early stop in the *clp-4* coding sequence and is a putative null. (**B**) Representative images of injured wild type and (**C**) *clp-4(-)* axons 24 hrs after injury. (**D**) *clp-4(-)* axons regenerate more frequently than wild type axons. but not in *parp(-)* mutant background. (**E**) Axon regeneration is not enhanced in the *parp(-); clp-4(-)* mutant relative to *parp(-)* or *clp-4(-)*, indicating the genes function in the same pathway to inhibit axon regeneration. (**F**). Loss of *clp-4* function does not increase the number of injured axons reach the dorsal cord (Full Regeneration), (**G**) nor does it change the ability of *parp(-)* axons to fully regenerate. Error bars indicate 95% confidence intervals. Fisher’s exact test was used to determine significance. *P < 0.05, **P < 0.01, ***P < 0.001, ****P < 0.0001.

Additional candidates included genes whose homologs interact with PARPs to regulate the DNA damage response *xpa-1*/XPA (Xeroderma pigmentosum comp grp A related) and *atm-1* (ATM ataxia telangectasia mutated family). Neither loss of function mutations in *xpa-1* or *atm-1* enhanced or suppressed axon regeneration in either wild-type or *parp(-)* animals (74.42% vs 68.97%, p>0.05) (Supplemental Figure 4A,B), suggesting neither gene regulates axon regeneration in coordination with the PARP genes. The role of PARPs in the immune response was also investigated by measuring axon regeneration in animals fed the *C. elegans* pathogen *P. aeruginosa* 14, which stimulates the immune response. While the immune response did stimulate axon regeneration, the interaction was not dependent on PARP function (Supplemental Figure 4C). Therefore, the immune response is unlikely to mediate the role of PARPs in axon regeneration.

### PARPs regulate synapse formation with *clp-4*

Our finding that PARPs inhibit both synapse reformation and axon regeneration suggests its downstream effectors may regulate one or both processes. We asked whether, like PARP genes, CLP-4 regulates both axon regeneration and synapse reformation after axon injury. We found that in the absence of *clp-4* function, the number of synaptic punctae in single axons that had fully regenerated to the dorsal cord were comparable to the number of synaptic puncta observed in single axons that had undergone mock surgery (18 vs 15, p>0.05) (Figure 5A). Therefore, synapse reformation is inhibited by *clp-4*. Because more synapses are formed in fully regenerated *clp-4(-)* axons compared to fully regenerated wild-type axons, these data indicate CLP-4, like the PARPs, specifically inhibits synapse formation independently from its role in axon regeneration.

**Figure 5.**
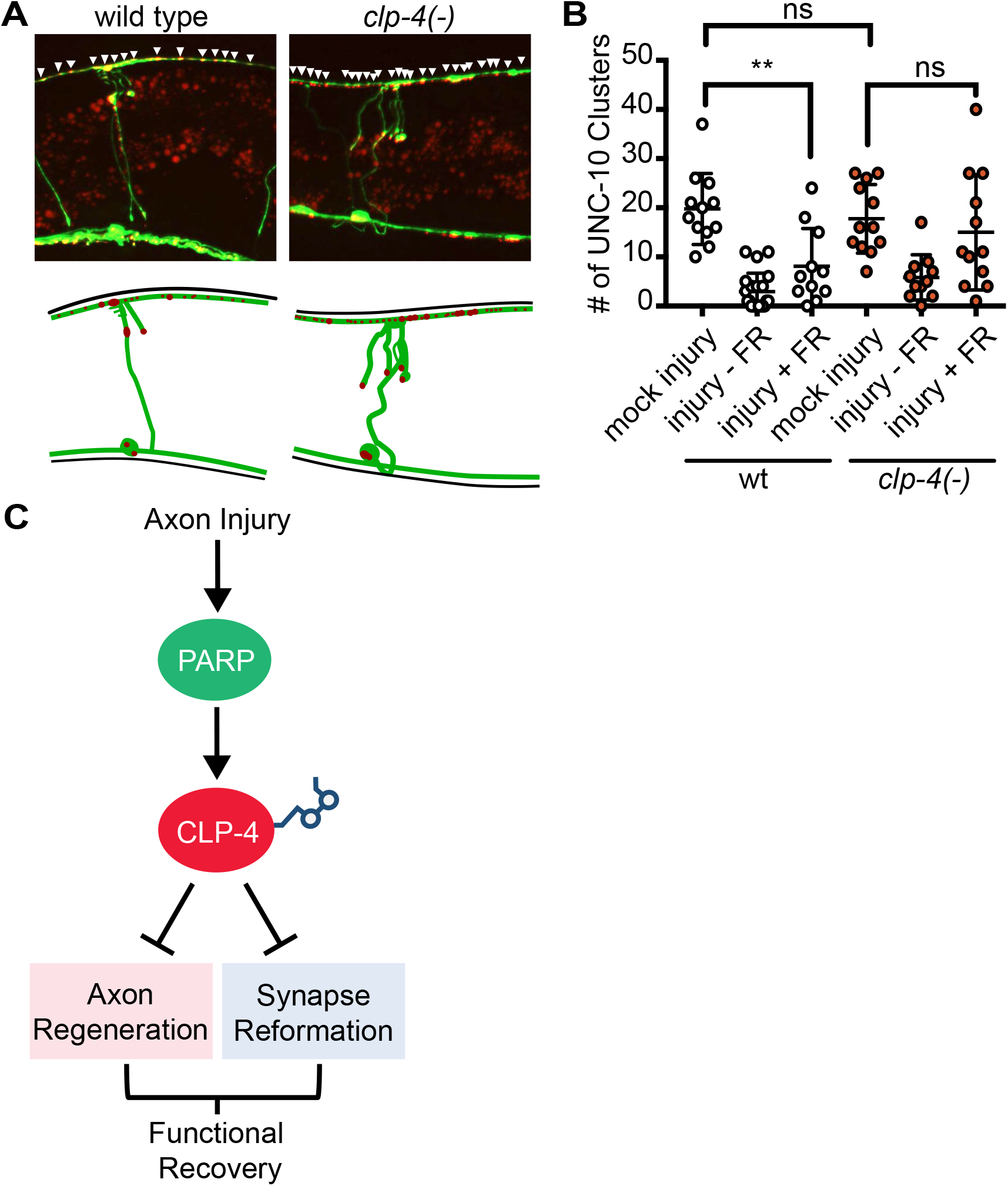
*clp-4* knockdown improves synapse reformation. (**A**) Representative images of synapse reformation in wild type and *clp-4(-)* mutants. Arrowheads indicate UNC-10::mCherry clusters. (**B**) Active zones degenerate and are reformed in *clp-4(-)* axons but are not reformed in wild type axons. Each dot represents one animal. Error bars represent standard deviation. Student’s t-test was used to determine significance, *P < 0.05, **P < 0.01, ***P < 0.001, ****P < 0.0001. (**C**) Proposed model of PARP mediated axon regeneration and synapse reformation. PARPs directly or indirectly regulate CLP-4 activity, which also inhibits both axon regeneration and synapse reformation, likely through separate downstream effectors. By enhancing both axon regeneration and synapse reformation, inhibiting PARP and CLP-4 function significantly improves the functional recovery of injured axons.

## Discussion

### PARP knockdown regulates both synapse reformation and axon regeneration

Here we find that poly (ADP-ribose) polymerases inhibit functional repair of the injured nervous system by regulating both synapse reformation and axon regeneration. Injury to the central nervous system or aged peripheral nervous system often results in a permanent loss of function, which persists because the injured axons do not regenerate. Moreover, multiple lines of evidence show that promoting axon regeneration is not sufficient to induce synaptic reformation and restore function (44–47). In parallel, epigenetic and transcriptional profiling has revealed that axons that are capable of regenerating adopt a transcriptional profile that resembles that of early developing axons prior to synaptogenesis, which includes a downregulation of synaptic gene expression (1, 46, 48). Therefore, by providing a novel understanding of how functional synapses can be rebuilt in a regenerating axon, our results provide a puzzle piece required for the restoration of full function after axon injury.

### A robust model to investigate functional synapse reformation in regenerated motor axons

*C. elegans* is a highly tractable model of axon regeneration and of synapse development. We combined novel and established regeneration and synapse assays to create an injury model that quantifies GABA motor axon regeneration, synapse formation, and function in wild type and *parp(-)* axons with single axon resolution. With this new model, we found that most wild-type GABA motor axons are incapable of reforming pre-injury levels of synaptic vesicles, active zone components, and functional interactions with postsynaptic muscle cells. A comparison of previous investigations reveals that *C. elegans* axons have cell-type specific outcomes to injury. The two most commonly investigated models of axon regeneration are injured mechanosensory neurons and motor neurons. Injured mechanosensory axons are capable of restoring their original functional interactions by fusing their severed axon fragments (49, 50). In the absence of fusion, partial restoration of function is dependent on accurate guidance of the regenerating axon to the post-synaptic cell (51). While injured motor neurons are incapable of axonal fusion, locomotor function is partially restored 24 hours after simultaneously severing a large number of GABA axons (11, 17, 52). Whether the partial restoration of function in these animals is caused by synaptic reformation or compensatory mechanisms was not determined in these assays. However, our finding that most wild type DD neurons have not reformed synapses at the time of these functional assays leaves the possibility that some functional restoration could indeed be caused by the additive effect of multiple regenerated axons that each have small numbers of synapses or could be attributed to compensatory mechanisms. In contrast to our findings in wild type motor axons, regenerated motor axons in mutant animals lacking RIC-7, an orphan protein that is required for mitochondrial transport and neurotransmitter release, regain pre-injury levels of synaptic vesicles in regenerated axons, partially regain active zone components, but recover limited function, in part because they aberrantly traffic synaptic vesicles to their dendrites (53). The differences in synapse dynamics reflect a distinction between *ric-7(-)* and wild type animals or between gabaergic and cholinergic axons. By establishing an understanding of whether injured wild type GABA motor axons are capable of functional regeneration, our data provides a fundamental baseline to investigate cell and genotype specific differences.

### DLK-1 regulates injury-induced synapse dynamics

We found that developmentally formed synapses degenerate in both wild type and *parp(-)* axons after injury. The MAPKKK *dlk-1*/DLK/Wnd is required for the decrease in synaptic vesicles in the distal stumps of axons that do not regenerate. Curiously, the distal stumps of severed motor axons are protected in severed *C. elegans* GABA axons, a characteristic that is inhibited by expression of *tir-1*/dSarm/SARM1, which regulates both axon degeneration and regeneration, and is dependent on *ric-7* (27, 54–56). Since the distal axon fragments are severed from their cell bodies, the difference in stability of synaptobrevin puncta between wild type and *dlk-1(-)* axons is not caused by defective trafficking from the cell body to the neuromuscular junction. Rather, our data is consistent with previous findings that activating DLK-1 or its homolog Wallenda in developing *C. elegans* and *D. melanogaster*, respectively, reduces expression of synaptic vesicles and active zone proteins and results in defective synaptic function (20, 28–31). DLK-1/DLK/LZK drives regeneration in *C. elegans*, *D. melanogaster*, and mammals. In *dlk-1(-)* animals, injured axons cannot initiate axon regeneration, and therefore we could not determine whether *dlk-1* regulates synapse reformation. An emerging theme in axons that are capable of regenerating is that synapse destabilization improves axon growth and that developmental synapses need to be cleared for new synapses to form (1, 46). These data suggest that manipulating DLK-1/DLK/LZK/Wnd expression might be a promising approach to both promote axon regeneration and synapse reformation across species.

### PARPs inhibit synapse reformation

This study reveals that PARPs inhibit functional recovery by regulating the reformation of new synaptic vesicles, active zones, and functional interactions between regenerating axons and their target cells. In *parp(-)* animals, we observed a similar amount of synaptic degeneration at the developmental neuromuscular junction compared to wild type animals. These data indicate the increased number of synaptic vesicles, active zones, and trans-synaptic interactions is caused by an increase in synapse reformation and not by a decrease in synapse destabilization in *parp(-)* animals. Nor is the increase in synapse number a consequence of axonal fusion, since the severed distal stump of each regenerated axon was visible at the time of synapse analysis. Of note, the number of synaptic vesicles and active zone proteins are significantly increased in individual regenerated *parp(-)* axons compared to individual regenerated wild type axons. Because both experiments were conducted on individual axons that had reached the dorsal cord, PARP function in synapse reformation is separable from its role in axon regeneration. Therefore, the PARPs likely inhibit each process in coordination with shared and divergent upstream activators and downstream effectors. The number of reformed synaptic vesicles in both wild type and *parp(-)* axons was variable and in many cases exceeded the number of synaptic vesicles in the same axons before injury. In contrast, the number and variability of UNC-10/RIM puncta match pre-injury axons, indicating the active zones may be a better representation of functional synapses. Knowing whether the new synapses are edited or continue to form with increased time will be critical to determining whether they are stable and whether PARPs also regulate synapse maintenance. Whether PARPs function in muscle cells to regulate the post-synapse in coordination with the pre-synapse is a particularly intriguing future direction.

Determining whether PARPs regulate synapse formation in mammals is made difficult by the complexity of the mammalian system. Targeting PARP-1 or PARP-9 by siRNA is sufficient to induce axon regeneration in injured primary cortical neurons in cell culture (11). However, while PARPs inhibit mammalian axon regeneration in vitro, chemical PARP inhibition did not restore axon regeneration to injured optic nerves *in vivo* (16). It is possible that within an *in vivo* context, strong inhibition of only one of the 18 PARPs is not sufficient to overcome parallel intrinsic and extrinsic inhibitory cues, including PTEN, chondroitin sulphate proteoglycans, and myelin associated inhibitors. Simultaneously blocking the function of multiple growth inhibitors improves the ability of axons to regenerate significantly more than knocking down the same inhibitors individually (7, 57). Whether inhibiting PARP function in combination with manipulations that reduce the extrinsic inhibition of axon regeneration will enhance axon regeneration and functional synapse formation will be critical to determining whether chemical PARP inhibition is a viable therapeutic approach to stimulate functional axon regeneration.

### PARPs regulate axon regeneration by PARylating substrates

Two lines of evidence suggest PARPs inhibit axon regeneration directly by PARylating substrate(s), and not by affecting the amount of available NAD^+^. First, the inhibitory effect of NAD^+^ on axon regeneration was abolished when PARPs were mutated. These data indicate that increased NAD^+^ inhibits axon regeneration by promoting the enzymatic activity of PARPs. Second, we previously found that PARGs, which remove poly (ADP-ribose) from substrates, have the opposite role in axon regeneration. The opposing effects on axon regeneration between PARPs and PARGs, suggests their contrasting enzymatic activities regulate axon regeneration. Interestingly, PARG expression is regulated by DLK-1 and is required for both DLK-1 induced axon regeneration and developmental synapse destabilization (11). Whether the enzymatic activity of PARGs also regulates synapse destabilization and reformation after injury remains to be determined.

### PARPs inhibit functional axon regeneration with CLP-4

We found that the CLP-4 calpain functions in coordination with PARPs to regulate both growth cone formation and synaptic reformation. These functions are consistent with known functions of calpains, which are calcium-dependent proteolytic enzymes implicated in developmental axonal outgrowth and synaptic pruning. For example, in *Xenopus laevis*, calpain activation by Ca^+^ influx through a TRPC1 channel reduces developmental axon outgrowth (58). Activated calpains also regulate synaptic plasticity during memory consolidation and developmental pruning (58–62). However, hyper-activation of calpain-2 can also contribute to detrimental memory loss in Alzheimer disease mouse model (63). In contrast, we found that *clp-4(-)* axons and synapses develop normally, suggesting *clp-4* is temporally regulated such that it functions specifically in injury-induced axon regeneration or that it functions redundantly with one of the other *C. elegans* calpain homologs during development. A more detailed molecular and cellular characterization of CLP-4 is required to understand how it interacts with the cytoskeleton and synaptic proteins in injured axons.

The absence of an additive regeneration phenotype in *parp(-);clp-4(-)* animals relative to *parp(-)* or *clp-4(-)* animals indicates CLP-4 functions with PARPs to regulate axon regeneration. Whether PARPs function upstream or downstream of calpain is not known, but both interactions are possible. In support of PARPs functioning upstream: 1) PARPs are known to PARylate calpains upon cellular stress (64, 65), 2) While *parp(-)* mutants display an increase in both growth cone initiation and extension, loss of *clp-4* function only affects growth cone formation. The stronger *parp(-)* phenotype suggests that PARPs likely function upstream to regulate multiple effectors and temporally fine-tune at least three components of functional axon regeneration: growth cone formation, axon elongation, and synaptic reformation, and 3) Calpains cleave synaptic proteins, which is consistent with the finding that they directly inhibit synapse reformation. However, in support of Calpains functioning upstream of PARPs is the finding that Calpains have also been shown to cleave PARPs. Considering the existing literature in sum, our data more strongly supports a model where PARPs regulate CLP-4 directly or indirectly via PARylation, activated CLP-4 in turn inhibits synapse reformation by cleaving synaptic components and ultimately achieves homeostasis by cleaving the PARPs. Future genetic and biochemical experiments will determine the nature of the relationship between the PARPs and CLP-4.

Together, these findings provide significant insight into the molecular mechanisms that regulate functional regeneration at both the level of regenerating axons and newly forming synapses. They highlight that functional axon regeneration is the cumulation of multiple processes, including injury sensing, growth cone formation, axon guidance, target recognition, synapse formation, and presumably, synapse pruning and maintenance. Each of these processes are intricately regulated in space and time during development and must be co-opted post developmentally in injured axons. Our finding that PARP knockdown restores function to individual axons by releasing inhibition of both axon regeneration and synapse formation suggests it is a relatively master regulator of the injury response. It remains to be determined whether disrupting multipotent genes such as PARPs or combinatorically manipulating individual components of the injury response will be required to effectively and specifically induce functional axon regeneration in a therapeutic setting. However, approaches that consider both axon regeneration and synapse reformation will be critical to restoring function to the injured nervous system.

## Acknowledgments

We thank all members of the Byrne laboratory, Dr. Alejandro Vasquez Rifo, Dr. Michael Francis, and Dr. Vivian Budnik for their experimental suggestions, insight and critical reading of the manuscript. We thank the *C. elegans* Gene Knockout Consortium (Mitani laboratory and Oklahoma Gene Knockout Consortium), the Caenorhabditis Genetics Center (CGC), which is funded by the NIH Office of Research Infrastructure Program (P40 OD010440), for providing strains and WormBase. We thank the Francis (UMass Chan Medical School), Hanna-Rose (Penn State), Ambros (UMass Chan Medical School) laboratories for reagents and expertise.

## Author contributions

M.Y.B contributed to conceptualization, investigation, interpretation and manuscript preparation. W.H contributed to investigation, interpretation and manuscript preparation. J.T.F. contributed to investigation and interpretation. M.J.A. contributed to interpretation and manuscript preparation. A.B.B contributed to conceptualization, interpretation and manuscript preparation.

## Data and materials availability

All data is available in the main text or the supplementary materials. Strains will be made available upon request.

## Materials and Methods

### Strains

Animals were maintained at 20°C on NGM plates containing OP50 *E. coli* according to standard methods. EG1285 *oxIs12[unc-47p::GFP + lin-15(+)] X,* NY2037 *ynIs37[flp- 13p::GFP] III,* ABC18 *wpIs39[unc-47p::mCherry] X,* ABC13 *ynIs37[flp-13p::GFP] III; wpIs39[unc-47p::mCherry] X,* CZ333 *juIs1[unc-25p::snb-1::GFP + lin-15(+)] IV,* ZM2246 *hpIs88[unc-25p::mCherry::unc-10 + lin-15(+)],* ABC60 *parp-1(ok988), parp-2(ok344); oxIs12 [unc-47p::GFP, lin-15(+)] X,* ABC35 *clp-4(o2808); oxIs12[unc-47p::GFP + lin- 15(+)] X,* ABC81 *tank-1(ok446); oxIs12[unc-47p::GFP + lin-15(+)] X,* ABC97 *xpa-1(ok698); oxIs12[unc-47p::GFP + lin-15(+)] X,* ABC118 *atm-1(gk186); oxIs12[unc- 47p::GFP + lin-15(+)] X,* ABC108 *pnc-1(pk9605); oxIs12[unc-47p::GFP + lin-15(+)] X,* ABC146 *parp-1(ok988); parp-2(ok344); pnc-1(pk9605); oxIs12[unc-47p::GFP + lin- 15(+)] X,* ABC113 *parp-1(ok988); parp-2(ok344); clp-4(o2808); oxIs12[unc-47p::GFP + lin-15(+)] X,* ABC27 *parp-1(ok988); parp-2(ok344)*; *ynIs37[flp-13p::GFP] III,* ABC61 *wpIs39[unc-47p::mCherry]; juIs1[unc-25p::snb-1::GFP + lin-15(+)] X,* ABC15 *parp- 1(ok988); parp-2(ok344); wpIs39[unc-47p::mCherry] X; juIs1[unc-25p::snb-1::GFP + lin- 15(+)] IV,* ABC159 *dlk-1(ju476); wpIs39[unc-47p::mCherry] X; juIs1[unc-25p::snb- 1::GFP + lin-15(+)] IV,* ABC96 *hpIs88[unc-25p::mCherry::unc-10 + lin-15+]; oxIs12[unc- 47p::GFP + lin-15(+)] X,* ABC153 *parp-1(ok988); parp-2(ok344); hpIs88[unc- 25p::mCherry::unc-10 + lin-15(+)]; oxIs12[unc-47p::GFP + lin-15(+)] X,* ABC144 *clp- 4(o2808); hpIs88[unc-25p::mCherry::unc-10 + lin-15+]; oxIs12[unc-47p::GFP + lin- 15(+)] X, pnc-1(pk9605)*.

### Axotomy and Microscopy

Axotomy experiments were performed as described in Byrne et al., 2011. Images were acquired with an Andor Zyla sCMOS camera on a Nikon NI-SSR-930959 microscope and NIS-Elements AR5.02.00 software, or a Perkin Elmer Precisely UltraVIEW VoX confocal imaging system mounted on a Zeiss AXIO (imager.M2) microscope and Volocity 6.3 software. Error bars indicate 95% confidence intervals. Significance was determined with a Fisher’s exact test.

### Functional Recovery

Functional locomotor recovery was assayed by specifically axotomizing DD2, DD3, DD4, and DD5 axons in *ynIs37* animals and quantifying dorsally-directed locomotion bias 24 hours later. Locomotion bias was quantified using a modified version of the assay described in (26). An individual worm’s movement was recorded with a MultiWorm Tracker v1.3.0 (https://sourceforge.net/projects/mwt/) (66)) for 300 seconds, 24-30 hours after injury. Worm tracks were analyzed by customized MATLAB scripts created by the Alkema lab. Dorsally-directed locomotion bias was quantified by averaging the radius of curvature for each 10 second segment of a single worm’s track over the 300 second recording. To calculate radius of curvature, a circle was fit to the track segment (*Circle Fit (Taubin method)*, Nikolai Chernov, MATLAB Central File Exchange) and the reciprocal of the radius of this circle was reported (1/r). The dorsal or ventral bias of the curve was scored manually using playback of the tracking results. Dorsal curves were marked as positive values and ventral curves as negative. For each experimental group animals that had fully regenerated 1 or 2 axons were scored. The average locomotion bias of 7-8 animals is reported. Error bars indicate standard error of the mean. Significance was determined with t-test.

### RNAi

RNAi clone (II-6J06) from the *C. elegans* RNAi feeding library was used to knock down *parp-2* (Kamath 2003). RNAi clone pMBP1 (*parp-1* RNAi) was constructed by cloning the *parp-1* open reading frame from mixed stage N2 cDNA into the RNAi feeding vector L4440 using primers TAGCGGCCGCGTCTTTTGTGGCACGGATCAGG and TAGCGGCCGCCATGTAAGTGAGGCCCAGTGGA. RNAi strains were cultured overnight in LB with 25 μg/mL carbenicillin and 12.5 μg/mL tetracycline. The resulting culture was diluted 1:100 in LB + 25 μg/mL carbenicillin for 6h (Jedrusik & Shulze, 2004) and 500µl RNAi culture was seeded on 95% NGM plates containing 25 μg/mL carbenicillin and 1mM IPTG. RNAi were fed to XE1375 *wpIs36[unc-47p::mCherry] I; wpSi1[unc-47p::rde-1::SL2::sid-1 + Cbr-unc-119(+)] II; eri-1(mg366) IV; rde-1(ne219) V; lin-15B(n744)* animals (34). The negative control L4440 RNAi was cultured and fed to XE1375 animals in parallel.

### Synapse Reformation

DD GABA motor neuron commissures were visualized using the DD specific *flp- 13::GFP* marker. The DD6 commissure was identified by analyzing the overlapping signal between *flp-13::GFP* expression and GABA specific *unc-47p::mCherry* expression (Supplemental Figure 1). Synaptic markers *juIs1[unc-25p::snb-1::GFP + lin- 15+]* and *hpIs88[unc-25p::mCherry::unc-10 + lin-15+]* were imaged 24 hours after axotomy with a ZEISS LSM700 imaging system mounted on ZEISS AXIO (Imager.Z2) microscope using Zen 2012 SP1 (black edition) software and a Perkin Elmer Precisely UltraVIEW VoX confocal imaging system mounted on a ZEISS AXIO (imager.M2) microscope using Volocity 6.3 software, respectively. Mean fluorescence and puncta number were quantified with Fiji (67). Error bars represent standard deviation. Significance was determined with Student’s t-test.

### Nicotinamide Riboside Supplementation

Nicotinamide Riboside (NR) (BOC sciences cas no 1341-23-7) was supplemented as described in (35). Animals were placed on plates containing 1.25 mM NR for 24 hours after injury.

### Pseudamonas aereuginosa Exposure

Slow killing (SK) plates seeded with *Pseudamonas aereuginosa* strains PA14 and *gacA* were prepared as described in (68). To activate the immune response post injury, animals were transferred from plates seeded with *E. coli* OP50 to plates seeded with the relevant *P. aereuginosa* 3 hours after axotomy. After 18h of infection, animals were recovered on OP50 plates for 3 hours. The 18h time point was chosen because a significant portion of animals die starting after 18 hours of exposure to PA14. Animals were mounted and regeneration was quantified 24h after axotomy.

---

**Figure S1.**
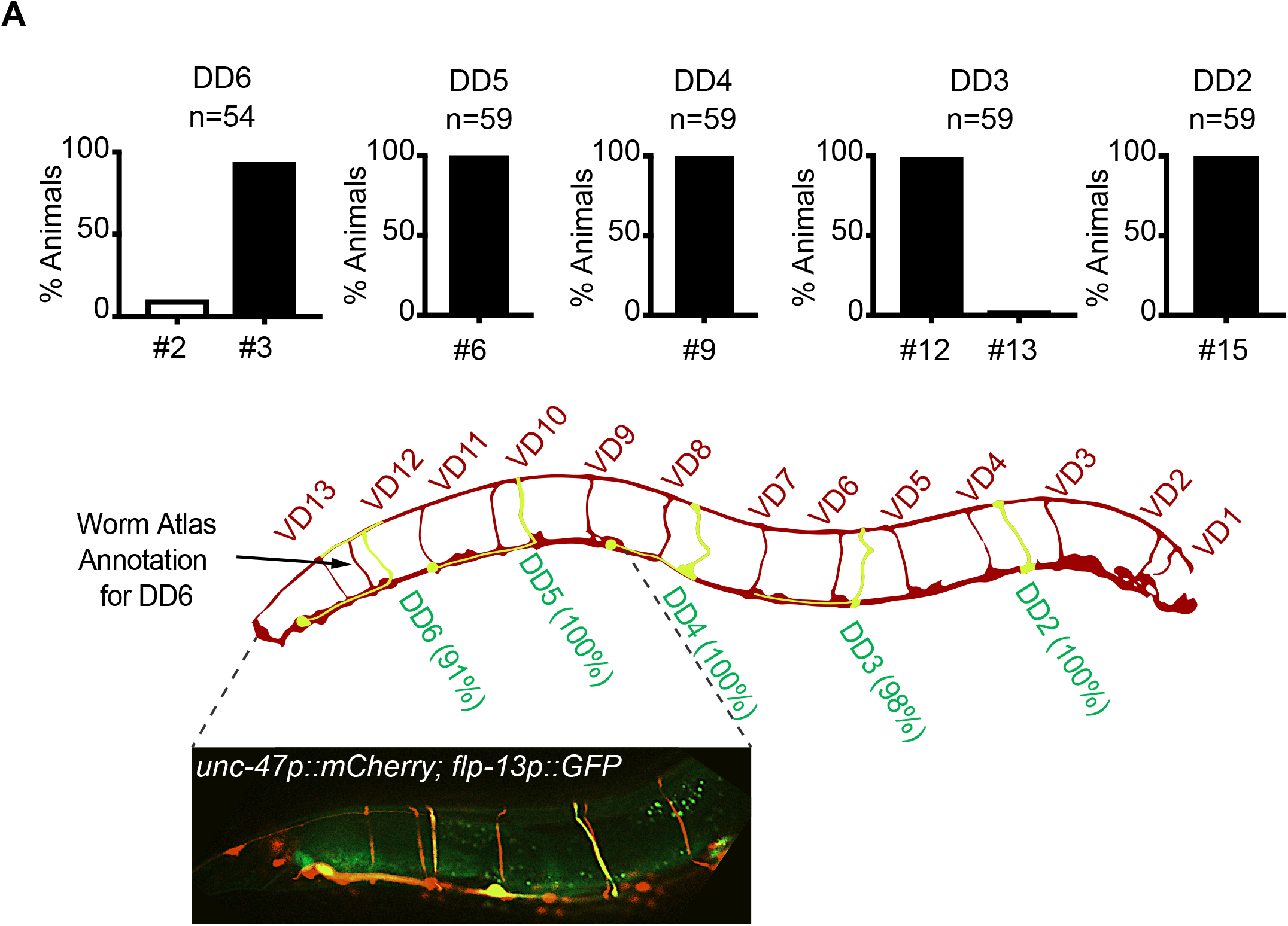
Commissure map of *C. elegans* DD and VD GABAergic neurons. (**A**) DD neurons are specifically labeled in green with *flp-13p::GFP* and all GABAergic neurons are labeled in red with *unc-47p::mCherry*. Commissure numbers start from the posterior end of the animal. DD axons were identified as those that co-express *flp-13p::GFP* and *unc-47p::mCherry*. Percentage indicates frequency of axon commissure number corresponding to respective DD neuron.

**Figure S2.**
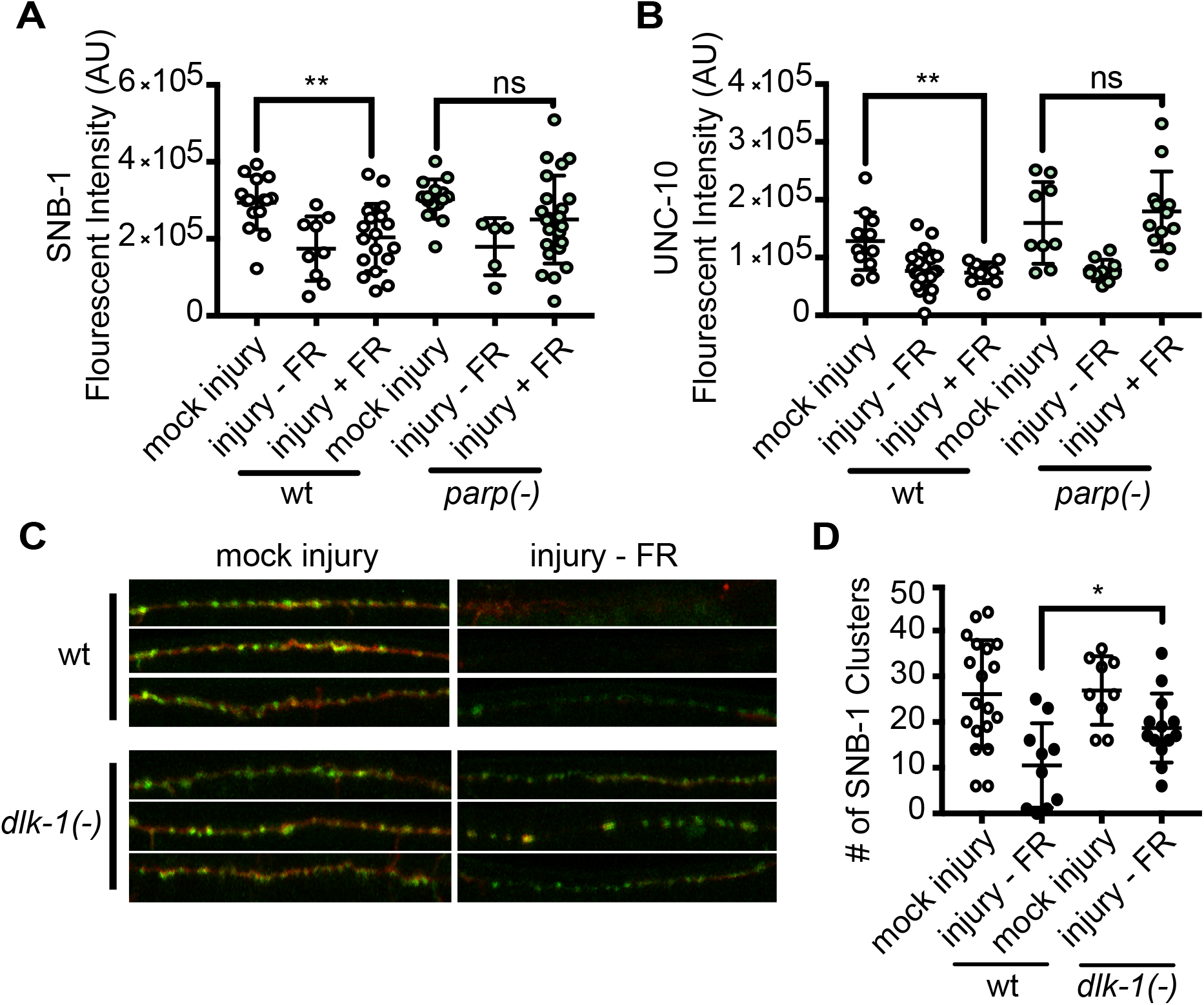
PARPs inhibit synapse reformation. (**A**) Quantification of synaptic SNB- 1::GFP fluorescent intensity in wild type animals and *parp(-)* mutants. (B) Quantification of synaptic UNC-10 fluorescent intensity in wild type animals and *parp(-)* mutants. (**C**) *dlk-1(-)* mutants have more SNB-1 puncta in injured axons compared to *wildtype* animals. Representative images show SNB-1::GFP clusters in dorsal cord axons of *wildtype* and *dlk-1* animals. Axons are labeled in red (*unc-47p::mCherry*) and SNB-1 puncta in green (*unc-25p::SNB-1::GFP*). For each genotype, mock injury and injury FR(-), were quantified. Quantification of SNB-1 clusters is shown in right panel. (**C**) Quantification of synaptic SNB-1 and UNC-10 fluorescent intensity in *wildtype* and *parp(-)* animals. (**D**) Dorsal cord regeneration phenotype in *wildtype* and *parp(-)* animals was quantified by measuring *unc-47p::mCherry* fluorescent intensity. Each dot represents an animal. Error bars represent standard deviation. t-test was used to determine significance. *P < 0.05, **P < 0.01, ***P < 0.001, ****P < 0.0001.

**Figure S3.**
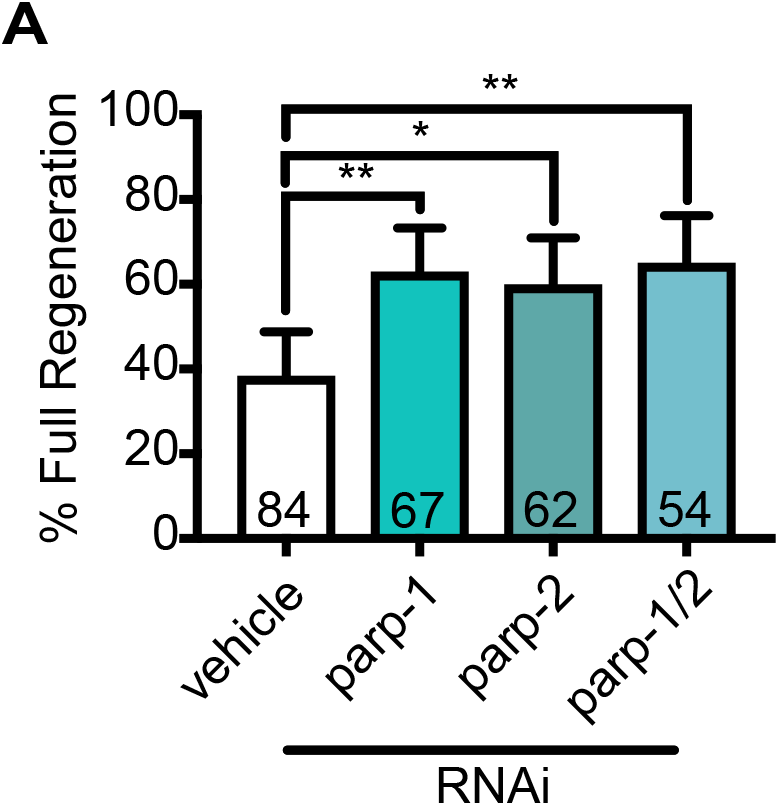
PARP genes regulate full regeneration cell-autonomously. (**A**) Full axon regeneration was quantified in animals in which RNAi is restricted to GABA neurons (see Methods). Animals were fed negative control L4440, *parp-1*, *parp-2*, or *parp-1 and parp-2* RNAi. RNAi targeting *parp-1*, *parp-2*, or both *parp-1* and *parp-2* increased axon regeneration relative to L4440. Error bars indicate 95% confidence intervals. Fisher’s exact test was used to determine significance. *P < 0.05, **P < 0.01, ***P < 0.001, ****P < 0.0001.

**Figure S4.**
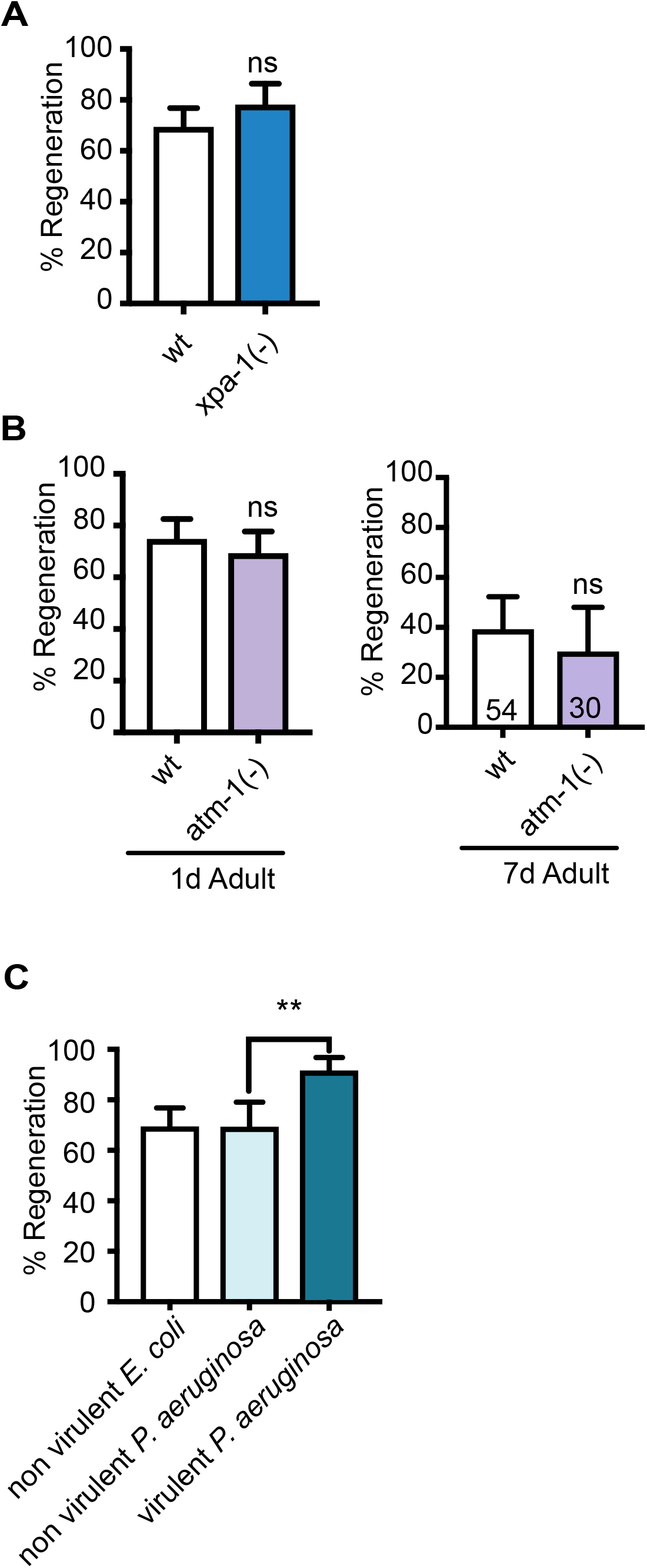
Candidate PARP interactors do not regulate axon regeneration in a PARP dependent manner. (**A**) Axon regeneration was measured in *wildtype* and *xpa- 1(-)* mutants. *xpa-1* has no effect on axon regeneration. (**B**) Axon regeneration was measured in *wildtype* and *atm-1(-)* mutants. *atm-1* has no effect on axon regeneration in 1 day adult or 7 day adult animals. (**C**) Exposure to virulent *P. aeruginosa* influences axon regeneration. (**D**) The increase in regeneration is not dependent on PARP function. (**E**) *parp(-)* mutant animals have no difference in survival rates compared to wild type animals in response to virulent *pseudomonas aeruginosa* strain, PA14. Survival curves of *wildtype* and *parp(-)* mutant animals exposed to PA14 starting at young adults are shown. Error bars indicate 95% confidence intervals. Fisher’s exact test was used to determine significance. *P < 0.05, **P < 0.01, ***P < 0.001, ****P < 0.0001.

**Figure S5.**
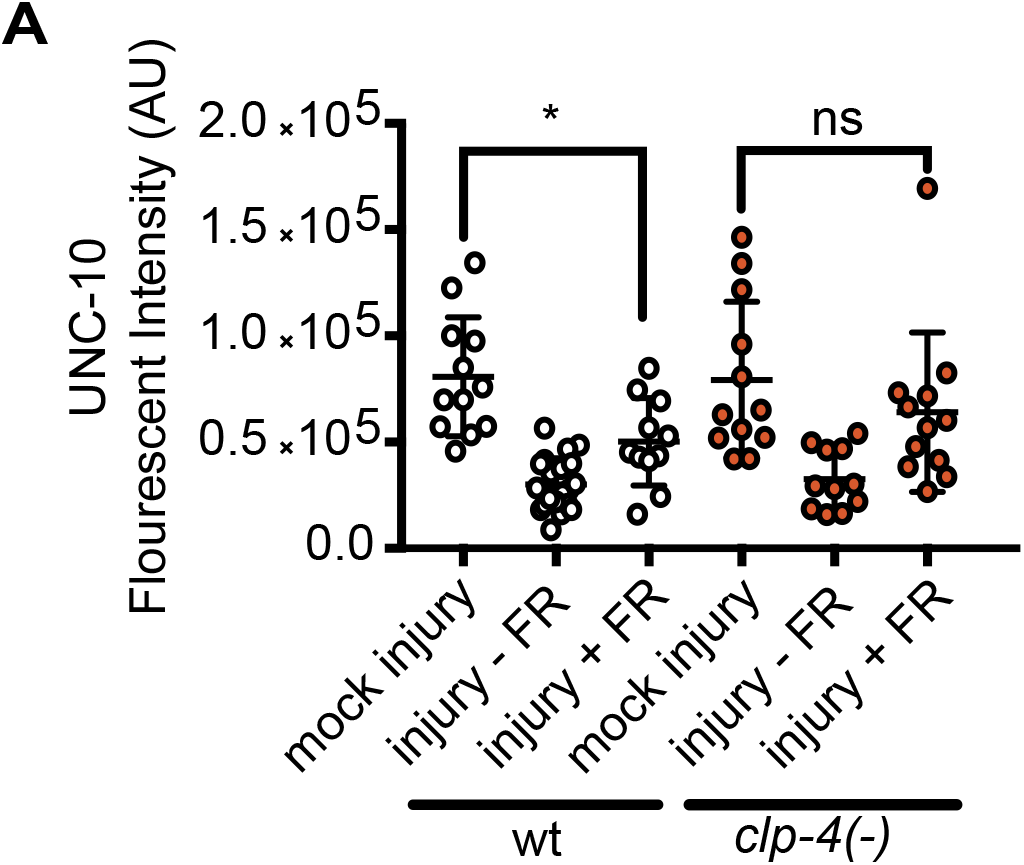
*clp-4* inhibits synapse reformation. (**A**) Quantification of synaptic UNC-10 fluorescent intensity in wild type animals and *clp-4(-)* mutants. For each genotype, mock injury, injury FR(-), and injury FR(+) were quantified. Each dot represents an animal. Error bars represent standard deviation. t-test was used to determine significance. *P < 0.05, **P < 0.01, ***P < 0.001, ****P < 0.0001.

## References

1. Mahar M, Cavalli V. Intrinsic mechanisms of neuronal axon regeneration. Nat Rev Neurosci. 2018;19(6):323–37.

2. Bradbury EJ, Burnside ER. Moving beyond the glial scar for spinal cord repair. Nat Commun. 2019;10(1):3879.

3. Fawcett JW. The Struggle to Make CNS Axons Regenerate: Why Has It Been so Difficult? Neurochem Res. 2020;45(1):144–58.

4. Byrne AB, Hammarlund M. Axon regeneration in *C. elegans*: Worming our way to mechanisms of axon regeneration. Exp Neurol. 2017;287(Pt 3):300–9.

5. Bei F, Lee HHC, Liu X, Gunner G, Jin H, Ma L, et al. Restoration of Visual Function by Enhancing Conduction in Regenerated Axons. Cell. 2016;164(1-2):219–32.

6. Liu Y, Wang X, Li W, Zhang Q, Li Y, Zhang Z, et al. A Sensitized IGF1 Treatment Restores Corticospinal Axon-Dependent Functions. Neuron. 2017;95(4):817–33 e4.

7. Wang Z, Mehra V, Simpson MT, Maunze B, Chakraborty A, Holan L, et al. KLF6 and STAT3 co-occupy regulatory DNA and functionally synergize to promote axon growth in CNS neurons. Sci Rep. 2018;8(1):12565.

8. Cafferty WB, McGee AW, Strittmatter SM. Axonal growth therapeutics: regeneration or sprouting or plasticity? Trends Neurosci. 2008;31(5):215–20.

9. Kaplan A, Bueno M, Hua L, Fournier AE. Maximizing functional axon repair in the injured central nervous system: Lessons from neuronal development. Dev Dyn. 2018;247(1):18–23.

10. Wood MD, Kemp SW, Weber C, Borschel GH, Gordon T. Outcome measures of peripheral nerve regeneration. Ann Anat. 2011;193(4):321–33.

11. Byrne AB, McWhirter RD, Sekine Y, Strittmatter SM, Miller DM, Hammarlund M. Inhibiting poly(ADP-ribosylation) improves axon regeneration. Elife. 2016;5.

12. Gupte R, Liu Z, Kraus WL. PARPs and ADP-ribosylation: recent advances linking molecular functions to biological outcomes. Genes Dev. 2017;31(2):101–26.

13. Brochier C, Jones JI, Willis DE, Langley B. Poly(ADP-ribose) polymerase 1 is a novel target to promote axonal regeneration. Proc Natl Acad Sci U S A. 2015;112(49):15220–5.

14. Ho TS-Y, Hees JT, Xu Z, Kawaguchi R, Biscola NP, Taub DG, et al. Screening for axon regeneration promoting compounds with human iPSC-derived motor neurons. bioRxiv. 2021:2021.11.02.466937.

15. Ame JC, Spenlehauer C, de Murcia G. The PARP superfamily. Bioessays. 2004;26(8):882–93.

16. Wang X, Sekine Y, Byrne AB, Cafferty WB, Hammarlund M, Strittmatter SM. Inhibition of Poly-ADP-Ribosylation Fails to Increase Axonal Regeneration or Improve Functional Recovery after Adult Mammalian CNS Injury. eNeuro. 2016;3(6).

17. Yanik MF, Cinar H, Cinar HN, Chisholm AD, Jin Y, Ben-Yakar A. Neurosurgery: functional regeneration after laser axotomy. Nature. 2004;432(7019):822.

18. Byrne AB, Edwards TJ, Hammarlund M. In vivo laser axotomy in *C. elegans*. J Vis Exp. 2011(51).

19. Hammarlund M, Nix P, Hauth L, Jorgensen EM, Bastiani M. Axon regeneration requires a conserved MAP kinase pathway. Science. 2009;323(5915):802-6.

20. Yan D, Wu Z, Chisholm AD, Jin Y. The DLK-1 kinase promotes mRNA stability and local translation in *C. elegans* synapses and axon regeneration. Cell. 2009;138(5):1005–18.

21. Byrne AB, Walradt T, Gardner KE, Hubbert A, Reinke V, Hammarlund M. Insulin/IGF1 signaling inhibits age-dependent axon regeneration. Neuron. 2014;81(3):561–73.

22. Watkins TA, Wang B, Huntwork-Rodriguez S, Yang J, Jiang Z, Eastham-Anderson J, et al. DLK initiates a transcriptional program that couples apoptotic and regenerative responses to axonal injury. Proc Natl Acad Sci U S A. 2013;110(10):4039–44.

23. Park KK, Liu K, Hu Y, Smith PD, Wang C, Cai B, et al. Promoting axon regeneration in the adult CNS by modulation of the PTEN/mTOR pathway. Science. 2008;322(5903):963–6.

24. Davis P, Zarowiecki M, Arnaboldi V, Becerra A, Cain S, Chan J, et al. WormBase in 2022-data, processes, and tools for analyzing *Caenorhabditis elegans*. Genetics. 2022;220(4).

25. E.M. Mslrrjskerhj. Identification and characterization of the vesicular GABA transporter. Nature. 1997;389(6653):870-6.

26. Donnelly JL, Clark CM, Leifer AM, Pirri JK, Haburcak M, Francis MM, et al. Monoaminergic orchestration of motor programs in a complex *C. elegans* behavior. PLoS Biol. 2013;11(4):e1001529.

27. Nichols ALA, Meelkop E, Linton C, Giordano-Santini R, Sullivan RK, Donato A, et al. The Apoptotic Engulfment Machinery Regulates Axonal Degeneration in *C. elegans* Neurons. Cell Rep. 2016;14(7):1673–83.

28. Nakata K, Abrams B, Grill B, Goncharov A, Huang X, Chisholm AD, et al. Regulation of a DLK-1 and p38 MAP kinase pathway by the ubiquitin ligase RPM-1 is required for presynaptic development. Cell. 2005;120(3):407–20.

29. Li J, Zhang YV, Asghari Adib E, Stanchev DT, Xiong X, Klinedinst S, et al. Restraint of presynaptic protein levels by Wnd/DLK signaling mediates synaptic defects associated with the kinesin-3 motor Unc-104. Elife. 2017;6:e24271.

30. Opperman KJ, Grill B. RPM-1 is localized to distinct subcellular compartments and regulates axon length in GABAergic motor neurons. Neural Dev. 2014;9:10.

31. Zhen M, Huang X, Bamber B, Jin Y. Regulation of presynaptic terminal organization by *C. elegans* RPM-1, a putative guanine nucleotide exchanger with a RING-H2 finger domain. Neuron. 2000;26(2):331–43.

32. Fire A, Xu S, Montgomery MK, Kostas SA, Driver SE, Mello CC. Potent and specific genetic interference by double-stranded RNA in *Caenorhabditis elegans*. Nature. 1998;391(6669):806-11.

33. Timmons L, Fire A. Specific interference by ingested dsRNA. Nature. 1998;395(6705):854.

34. Firnhaber C, Hammarlund M. Neuron-specific feeding RNAi in *C. elegans* and its use in a screen for essential genes required for GABA neuron function. PLoS Genet. 2013;9(11):e1003921.

35. Wang W, McReynolds MR, Goncalves JF, Shu M, Dhondt I, Braeckman BP, et al. Comparative Metabolomic Profiling Reveals That Dysregulated Glycolysis Stemming from Lack of Salvage NAD+ Biosynthesis Impairs Reproductive Development in *Caenorhabditis elegans*. J Biol Chem. 2015;290(43):26163–79.

36. Kim KW, Tang NH, Piggott CA, Andrusiak MG, Park S, Zhu M, et al. Expanded genetic screening in *Caenorhabditis elegans* identifies new regulators and an inhibitory role for NAD(+) in axon regeneration. Elife. 2018;7.

37. Erener S, Petrilli V, Kassner I, Minotti R, Castillo R, Santoro R, et al. Inflammasome-activated caspase 7 cleaves PARP1 to enhance the expression of a subset of NF-kappaB target genes. Mol Cell. 2012;46(2):200–11.

38. Haince JF, Kozlov S, Dawson VL, Dawson TM, Hendzel MJ, Lavin MF, et al. Ataxia telangiectasia mutated (ATM) signaling network is modulated by a novel poly(ADP-ribose)-dependent pathway in the early response to DNA-damaging agents. J Biol Chem. 2007;282(22):16441–53.

39. Knezevic CE, Wright G, Rix LLR, Kim W, Kuenzi BM, Luo Y, et al. Proteome-wide Profiling of Clinical PARP Inhibitors Reveals Compound-Specific Secondary Targets. Cell Chem Biol. 2016;23(12):1490–503.

40. Ko HL, Ren EC. Functional Aspects of PARP1 in DNA Repair and Transcription. Biomolecules. 2012;2(4):524–48.

41. Pleschke JM, Kleczkowska HE, Strohm M, Althaus FR. Poly(ADP-ribose) binds to specific domains in DNA damage checkpoint proteins. J Biol Chem. 2000;275(52):40974–80.

42. Wang Y, Kim NS, Li X, Greer PA, Koehler RC, Dawson VL, et al. Calpain activation is not required for AIF translocation in PARP-1-dependent cell death (parthanatos). J Neurochem. 2009;110(2):687–96.

43. Joyce PI, Satija R, Chen M, Kuwabara PE. The atypical calpains: evolutionary analyses and roles in *Caenorhabditis elegans* cellular degeneration. PLoS Genet. 2012;8(3):e1002602.

44. Carlin D, Golden JP, Mogha A, Samineni VK, Monk KR, Gereau RWt, et al. Deletion of Tsc2 in Nociceptors Reduces Target Innervation, Ion Channel Expression, and Sensitivity to Heat. eNeuro. 2018;5(2).

45. Carlin D, Halevi AE, Ewan EE, Moore AM, Cavalli V. Nociceptor Deletion of Tsc2 Enhances Axon Regeneration by Inducing a Conditioning Injury Response in Dorsal Root Ganglia. eNeuro. 2019;6(3).

46. Kiyoshi C, Tedeschi A. Axon growth and synaptic function: A balancing act for axonal regeneration and neuronal circuit formation in CNS trauma and disease. Dev Neurobiol. 2020;80(7-8):277–301.

47. Wang Z, Reynolds A, Kirry A, Nienhaus C, Blackmore MG. Overexpression of Sox11 promotes corticospinal tract regeneration after spinal injury while interfering with functional recovery. J Neurosci. 2015;35(7):3139–45.

48. Venkatesh I, Mehra V, Wang Z, Califf B, Blackmore MG. Developmental Chromatin Restriction of Pro-Growth Gene Networks Acts as an Epigenetic Barrier to Axon Regeneration in Cortical Neurons. Developmental Neurobiology. 2018;78(10):960–77.

49. Abay ZC, Wong MY, Teoh JS, Vijayaraghavan T, Hilliard MA, Neumann B. Phosphatidylserine save-me signals drive functional recovery of severed axons in *Caenorhabditis elegans*. Proc Natl Acad Sci U S A. 2017;114(47):E10196–E205.

50. Basu A, Dey S, Puri D, Das Saha N, Sabharwal V, Thyagarajan P, et al. let-7 miRNA controls CED-7 homotypic adhesion and EFF-1-mediated axonal self-fusion to restore touch sensation following injury. Proc Natl Acad Sci U S A. 2017;114(47):E10206–E15.

51. Basu A, Behera S, Bhardwaj S, Dey S, Ghosh-Roy A. Regulation of UNC-40/DCC and UNC-6/Netrin by DAF-16 promotes functional rewiring of the injured axon. Development. 2021;148(11):dev198044.

52. El Bejjani R, Hammarlund M. Notch signaling inhibits axon regeneration. Neuron. 2012;73(2):268–78.

53. Ding C, Hammarlund M. Aberrant information transfer interferes with functional axon regeneration. eLife. 2018;7:e38829.

54. Czech VL, O’Connor LC, Philippon B, Norman E, Byrne AB. TIR-1/SARM1 inhibits axon regeneration and promotes axon degeneration. eLife. 2023;12:e80856.

55. Rawson RL, Yam L, Weimer RM, Bend EG, Hartwieg E, Horvitz HR, et al. Axons degenerate in the absence of mitochondria in *C. elegans*. Curr Biol. 2014;24(7):760–5.

56. Hao Y, Hu Z, Sieburth D, Kaplan JM. RIC-7 promotes neuropeptide secretion. PLoS Genet. 2012;8(1):e1002464.

57. Jin D, Liu Y, Sun F, Wang X, Liu X, He Z. Restoration of skilled locomotion by sprouting corticospinal axons induced by co-deletion of PTEN and SOCS3. Nat Commun. 2015;6(1):8074.

58. Kerstein PC, Jacques-Fricke BT, Rengifo J, Mogen BJ, Williams JC, Gottlieb PA, et al. Mechanosensitive TRPC1 channels promote calpain proteolysis of talin to regulate spinal axon outgrowth. J Neurosci. 2013;33(1):273–85.

59. Khoutorsky A, Yanagiya A, Gkogkas CG, Fabian MR, Prager-Khoutorsky M, Cao R, et al. Control of synaptic plasticity and memory via suppression of poly(A)-binding protein. Neuron. 2013;78(2):298–311.

60. Nagayoshi T, Isoda K, Mamiya N, Kida S. Hippocampal calpain is required for the consolidation and reconsolidation but not extinction of contextual fear memory. Mol Brain. 2017;10(1):61.

61. Schuldiner O, Yaron A. Mechanisms of developmental neurite pruning. Cell Mol Life Sci. 2015;72(1):101–19.

62. Yaron A, Schuldiner O. Common and Divergent Mechanisms in Developmental Neuronal Remodeling and Dying Back Neurodegeneration. Curr Biol. 2016;26(13):R628–R39.

63. Ahmad F, Das D, Kommaddi RP, Diwakar L, Gowaikar R, Rupanagudi KV, et al. Isoform-specific hyperactivation of calpain-2 occurs presymptomatically at the synapse in Alzheimer’s disease mice and correlates with memory deficits in human subjects. Sci Rep. 2018;8(1):13119.

64. Jungmichel S, Rosenthal F, Altmeyer M, Lukas J, Hottiger MO, Nielsen ML. Proteome-wide identification of poly(ADP-Ribosyl)ation targets in different genotoxic stress responses. Mol Cell. 2013;52(2):272–85.

65. Zhang Y, Wang J, Ding M, Yu Y. Site-specific characterization of the Asp- and Glu-ADP-ribosylated proteome. Nat Methods. 2013;10(10):981–4.

66. Husson SJ, Costa WS, Schmitt C, Gottschalk A. Keeping track of worm trackers. WormBook. 2013:1–17.

67. Schindelin J, Arganda-Carreras I, Frise E, Kaynig V, Longair M, Pietzsch T, et al. Fiji: an open-source platform for biological-image analysis. Nat Methods. 2012;9(7):676–82.

68. Tan MW, Rahme LG, Sternberg JA, Tompkins RG, Ausubel FM. Pseudomonas aeruginosa killing of *Caenorhabditis elegans* used to identify *P. aeruginosa* virulence factors. Proc Natl Acad Sci U S A. 1999;96(5):2408–13.

